# Elongated cells drive morphogenesis in a surface-wrapped finite element model of germband retraction

**DOI:** 10.1101/504548

**Authors:** W. T. McCleery, J. Veldhuis, G. W. Brodland, M. E. Bennett, M. S. Hutson

## Abstract

During *Drosophila* embryogenesis, the germband first extends to curl around the posterior end of the embryo, and then retracts back; however, retraction is not simply the reversal of extension. At a tissue level, extension is coincident with ventral furrow formation, and at a cellular level, extension occurs via convergent cell neighbor exchanges in the germband while retraction involves only changes in cell shape. To understand how cell shapes, tissue organization and cellular forces drive germband retraction, we investigate this process using a whole-embryo, surface-wrapped cellular finite element model. This model represents two key epithelial tissues – amnioserosa and germband – as adjacent sheets of 2D cellular finite elements that are wrapped around an ellipsoidal 3D approximation of an embryo. The model reproduces the detailed kinematics of *in vivo* retraction by fitting just one free model parameter, the tension along germband cell interfaces; all other cellular forces are constrained to follow ratios inferred from experimental observations. With no additional parameter adjustments, the model also reproduces failures of retraction when amnioserosa cells are removed to mimic U-shaped mutants or laser-microsurgery experiments. Surprisingly, retraction in the model is robust to changes in cellular force values, but is critically dependent on starting from a configuration with highly elongated amnioserosa cells. Their extreme cellular elongation is established during the prior process of germband extension and is then used to drive retraction. The amnioserosa is the one tissue whose cellular morphogenesis is reversed in germband extension and retraction – serving as a store of morphological information that coordinates the forces needed to retract the germband back to its pre-extension position and shape. In this case, and perhaps more generally, cellular force strengths are less important than the carefully established cell shapes that direct them.

## INTRODUCTION

Development of an embryo or embryogenesis is a dynamic process involving organism-wide coordination of multiple cell and tissue types. Such coordination is a key feature of embryonic epithelia in which cells and tissues deform while tightly adhering to their neighbors. Coordinated cellular forces have been studied and modeled for several episodes of epithelial development in *Drosophila melanogaster* embryos, including ventral furrow invagination (4-12), germband extension (13-26), and dorsal closure (27-46). More recently, studies have begun to elucidate the cellular forces driving another major episode of *Drosophila* embryogenesis known as germband retraction (2, 47, 48). Prior work on the mechanics of retraction has drawn inferences from the stress fields within individual germband segments; however, to capture the coordinated mechanics of the entire process, one must consider cells and segments spanning the posterior-most three-quarters of the embryo surface. Here we present a whole-embryo, cellular finite element model that reproduces germband retraction, that elucidates how forces are coordinated across two key tissues – germband and amnioserosa – and that explores the robustness of retraction and its critical dependencies on cell shape and dynamic cellular forces.

Germband retraction occurs midway through *Drosophila* embryogenesis (Bownes stage 12), following germband extension and preceding dorsal closure. When retraction begins, the two key tissues form interlocking U-shapes, similar to the two-piece cover of a baseball (Fig. 1A and Movie S1 in Supplemental Material). The germband lies along the ventral and dorsal surfaces, bending around the embryo’s posterior end; the amnioserosa drapes over the embryo’s lateral flanks, bending across the anteriodorsal surface. Both tissues are cellular monolayers (47), but their cells are packed in the tissue plane with different shapes. Cells in the germband are roughly **isodiametric** regular polygons, while those in the amnioserosa are highly elongated (Fig. 1B, see Supplemental Material for a glossary of terms highlighted in bold). Over the next two hours, the germband retracts around the posterior end of the embryo to occupy the ventrolateral surface, and the amnioserosa contracts to a compact and roughly elliptical shape on the dorsal surface (Fig. 1A and B). During this process, germband cells become slightly elongated dorsolaterally and amnioserosa cells round up – processes that occur without significant cellular neighbor exchange and with a tight correlation between cell and tissue shape changes (47). Germband retraction fails in several known genotypes, most notably among the U-shaped mutants (3). Based on observations from these U-shaped mutants, from genetic disruptions of cell-matrix interactions, and from laser-microsurgery experiments, researchers have concluded that amnioserosa and germband cells have a mix of both active and passive roles (1-3, 47-52).

**FIGURE 1.**
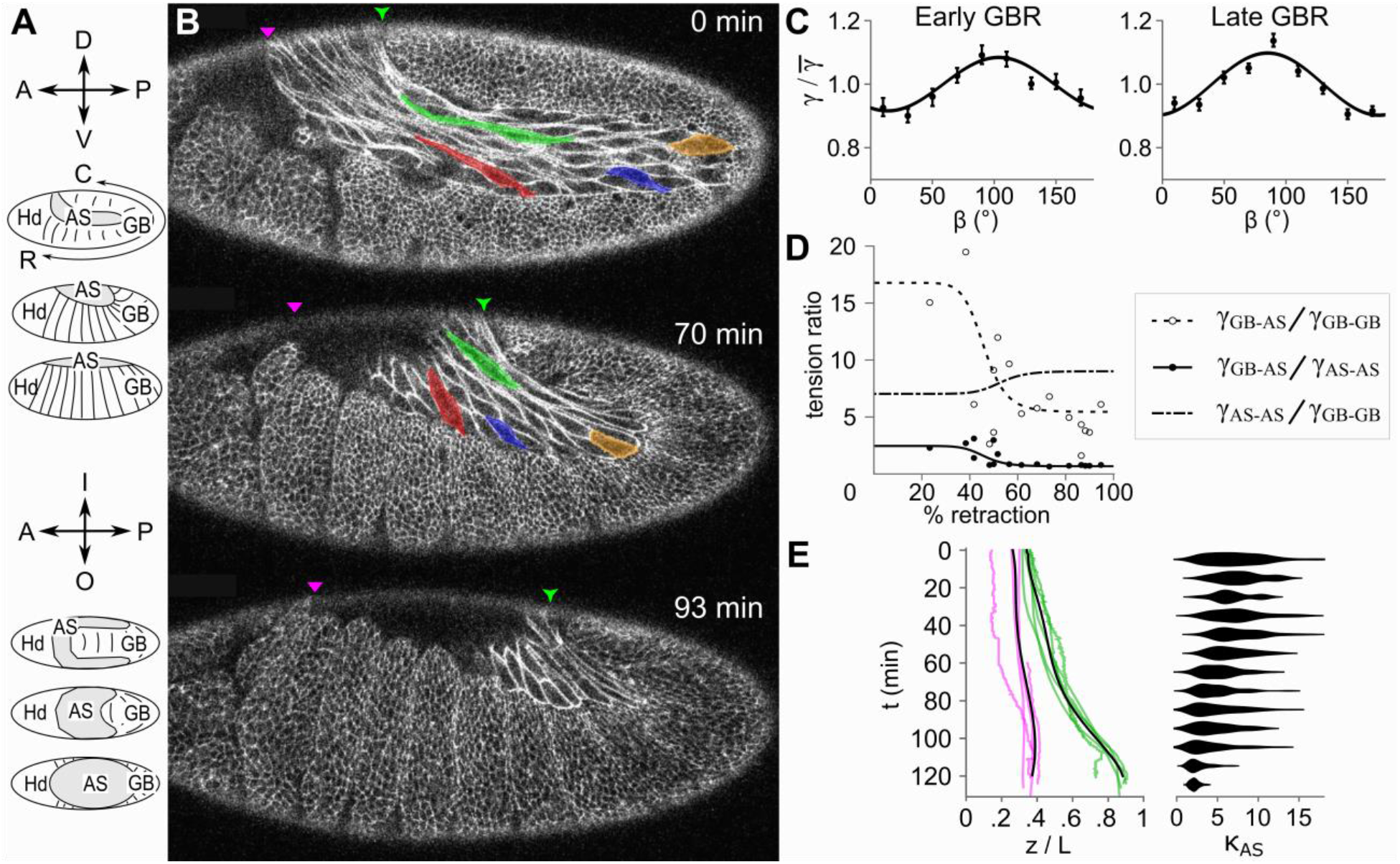
Measuring in vivo dynamics and relative tensions during germband retraction. (A) Schematic of germband retraction from lateral and dorsal views. Orientation axis labels: A, anterior; P, posterior; D, dorsal; V, ventral; R, rostral; C, caudal; I and O, lateral sides into and out of page. Tissue labels: Hd, head; AS, amnioserosa; GB, germband. (B) Time-lapse images from a lateral view of germband retraction in vivo. Anterior is left and dorsal up in this and all subsequent figures. Archicephalon and telson positions are respectively marked by magenta and green arrowheads. Morphology of select amnioserosa cells highlighted by color overlays. (C) Measurements of polarized germband cell-edge tensions as a function of orientation relative to the rostral-caudal axis. (D) Ratios of key cell-edge tensions determined from the shape of the GB-AS boundary. Symbols represent analysis of nine embryos imaged at different stages (% retraction). Solid and dashed lines are sigmoidal fits to the data; dot-dash line is a ratio of the other two fits. (E) Kinematic analysis of germband retraction (n = 5 embryos) showing time-dependent archicephalon (magenta) and telson (green) positions (z/L = position / embryo length). Smooth quantic regression lines overlain in black. Adjacent histograms show time-dependent distributions of amnioserosa cell aspect ratios (κ_AS_, n = 49-104 cells from 4 embryos).

The key tissue movements of germband retraction are clearly three-dimensional, but take place almost exclusively on the surface of the embryo. We thus model this process using a set of connected 2D epithelial cells that are wrapped over a rigid, 3D ellipsoidal core or **last**. This simplification reduces model run times while still capturing local and long-distance mechanical coordination. In this report, we first use experimental observations to determine or constrain model parameters. We then estimate remaining parameters by fitting model results to the observed **kinematics** of retraction. The resulting best-fit model accurately reproduces both normal germband retraction and its failure when amnioserosa mechanics are disrupted by mutation or microsurgery. We finally use the model to explore which aspects of cellular mechanics are critical. Surprisingly, retraction is robust to variations in cellular tensions: fourfold changes in any of the tensions result in at least partial retraction, albeit with altered kinematics. Retraction does fail however without the initial, highly elongated shapes of amnioserosa cells. These cell shapes are taken as initial conditions in the model, but they are determined in the embryo by cell and tissue movements in the previous morphogenetic process. The model is thus able to reveal a key and previously unappreciated link between germband extension and retraction. These processes are not the reverse of one another, but the second is clearly contingent on the cell geometry and topological connectedness achieved during the first. Such contingency may well be a ubiquitous aspect of embryonic development.

## MATERIALS AND METHODS

### Imaging and cell analysis

Live *Drosophila* embryos were imaged on a laser scanning confocal microscope (Axiovert-135TV/LSM-410; Carl Zeiss, Thornwood, NY) using a 40x, 1.3NA oil-immersion objective. Germband retraction was tracked in ubi-DE-Cad-GFP flies (Drosophila Genetic Research Center, Kyoto, Japan), which ubiquitously express E-Cadherin-GFP to label adherens junctions between cells (53). Additional whole-embryo images collected via Selective Plane Illumination Microscopy (SPIM) were used from previously published sources (supplementary videos 2, 3 and 5 from 54, supplementary video 4 from 55). Measurements of cell shapes and movements were performed using FIJI (56), Seedwater Segmenter for watershed-based segmentation (57), and CellFIT for inference of cellular tensions (58). Subsequent analysis and construction of plots was conducted in Mathematica 10 (Wolfram Inc., Champaign, IL).

### Computational modeling

Our model of germband retraction represents three epithelial tissues – amnioserosa (AS), germband (GB) and head (Hd) – as connected meshes of 2D cells wrapped over the surface of a 3D prolate ellipsoid (axes ratio of 1:1:2.5, which approximates the shape of a *Drosophila* embryo as shown in Fig. 2). The model does not include interior cells, such as precursors to the midgut, central nervous system, and mesoderm, because these cells have not yet formed cohesive tissues through which forces could be propagated (47 and others cited therein). Each surface cell is represented by a cellular finite element that is a polygon of fixed area, with edges that carry a defined active tension, and with vertices connected by two orthogonal sets of dashpots (59, 60). The fixed area represents incompressibility of both cytoplasm and the in-plane apical cytoskeleton (59). The active cortical tensions represent a combination of actomyosin contractions and the tangential equivalent of cell-cell adhesion (59, 61); and the dashpot networks damp changes in cell shape due to the effective viscosity of cytoplasmic flow and cytoskeletal remodeling (59, 62). The model is an idealization that does not explicitly include forces perpendicular to cell edges, e.g., pressures or contractions due to apical-medial actomyosin. Pressures are implicit in the cellular area constraints and apical-medial foci of actomyosin are not as prevalent during retraction as they are during preceding germband extension (21) or subsequent dorsal closure (38). All cells are modeled using the same effective viscosity, but the active tensions vary depending on the pair of cell types meeting at each edge and potentially on the edge orientation. The latter is applicable to germband cells for which there is evidence of planar cell polarity (15, 63 and results below).

**FIGURE 2.**
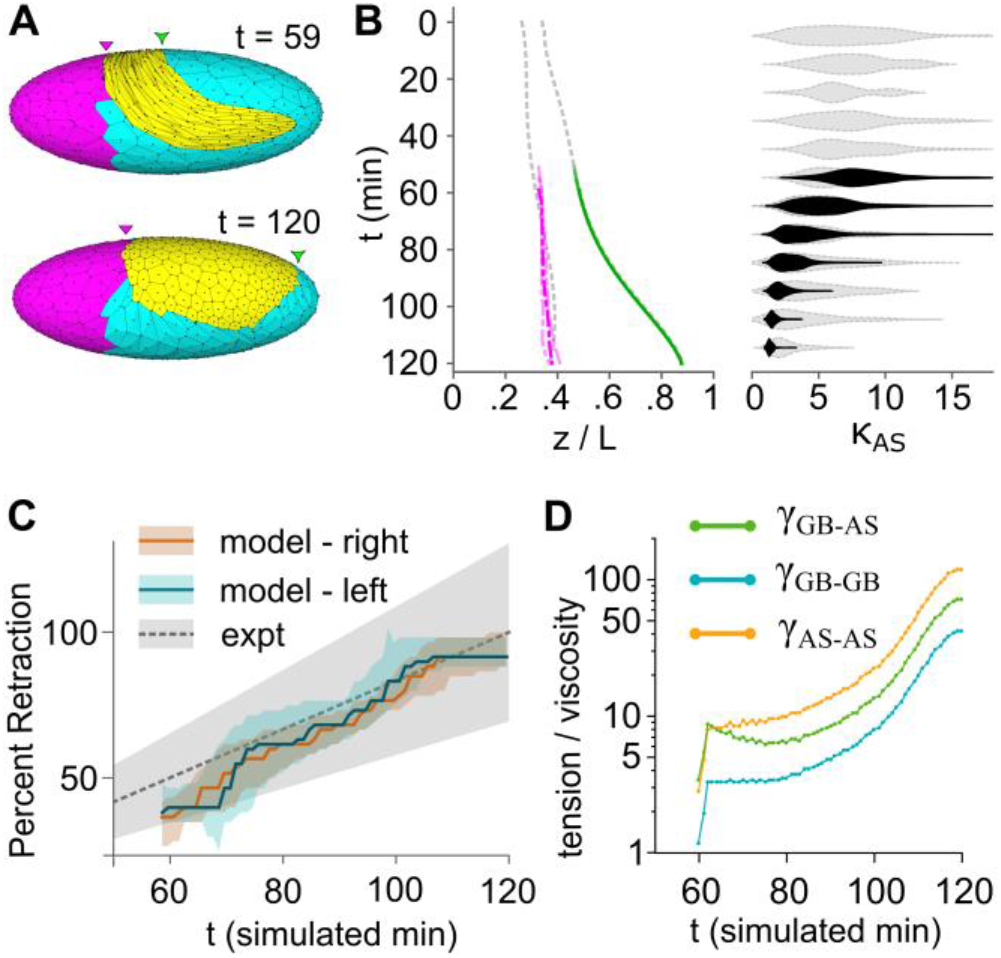
Best-fit model matched to *in vivo* kinematics. (A) Initial and final morphologies of the best-fit model. (B) Comparison of archicephalon and telson movements: best-fit models (dot-dashed magenta and solid green respectively) versus *in vivo* data (dashed gray; smooth quantic regressions shown). Comparison of amnioserosa cell aspect ratio distributions: best-fit model (black) versus *in vivo* data (light gray). (C) Comparative staging based on shape of the GB-AS boundary. Shaded gray area for experimental data is from the *in vivo* standard of Lynch et al (2). (D) Tension-viscosity ratios for the best-fit model. Plot indicates actual values used in simulation γ_GB-GB_ should be scaled by the square root of 10 to compare to *in vivo* tensions. Panels A, C and D show results from a best-fit using one of three initial cell meshes; panel B shows best-fit results for all three meshes. Color coding for similar plots applies to all subsequent figures.

The finite element engine assumes **low Reynolds number** conditions (64) and incrementally displaces each vertex along the ellipsoidal surface according to the governing equation,

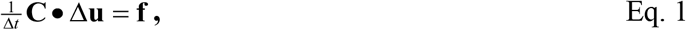

where Δ*t* is a simulation time step, **C** is a damping matrix resulting from the current geometry of the viscous dashpot networks, **Δu** is a vector of incremental node displacements, and **f** is a vector of the imbalanced active tensions applied to each node. This system of equations is augmented by applicable constraints (surface confinement and constant cell area) implemented via Lagrange multipliers. Cell intercalation does not occur during germband retraction (2, 47), so the model does not permit cellular neighbor exchanges. The model was custom-built based on previously published finite element engines (60, 65, 66) and adapted to handle a constraining ellipsoidal last. Initial cell configurations were designed and produced in Mathematica 10. Model results were visualized through a custom-built simulation viewer, ChiChi3D.

## RESULTS

Within the structure defined above, even the simplest time-independent model has nine free parameters: active edge tensions (*γ*) for six types of cell-cell interfaces; a direction (*β*_0_) and magnitude (*α*) for tension anisotropy in polarized germband cells; and an effective viscosity (*η*) for all cells. Since none of these have been previously measured experimentally, we first present *in vivo* measurements that directly determine some parameters, constrain relationships among others, and provide metrics to which the model and its remaining free parameters can be fit.

### Measuring germband polarization

Previous studies suggest that germband cells are polarized in plane (15, 48, 63). To measure the mechanical properties of this polarization, we imaged fly embryos expressing E-cadherin-GFP, segmented the images to determine cell boundaries and triple-junction angles throughout the germband (57), and subjected this geometry to cellular force inference to estimate the relative tension along each cell-cell interface (58). For early germband retraction, we imaged 7 embryos, segmented 71 cell patches containing 6,118 cells, and were able to estimate tensions for 11,399 cell edges. For late germband retraction, the numbers were 10 embryos, 104 segmented patches, 8,073 cells, and 16,211 cell edges. Results of this analysis are summarized in Fig. 1C: germband cell edges oriented perpendicular to the **rostral-caudal axis** are ∼10% stronger than the mean; those oriented parallel are ∼10% weaker; and this conclusion holds throughout retraction. The early and late anisotropies can be fit to a simple functional form,

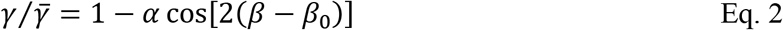

to respectively yield *α* = 0.08 ± 0.01, *β*_0_ = 15º ± 4º and *α* = 0.10 ± 0.01, *β*_0_ = −5º ± 4º. Interestingly, in both cases, the largest cortical tensions are oriented towards the amnioserosa (*β* ∼ 90º), which is parallel to the direction in which germband cells elongate. Germband cell tensions thus resist rather than drive their own elongation.

### Measuring active tensions for classes of cell-cell interfaces

Having directly determined the direction and magnitude of tension anisotropy in the germband, we then conducted additional force inference to constrain relationships among the three key types of active tensions: AS-AS, GB-AS and GB-GB. To do so, we imaged the GB-AS boundary near the crook of the germband. As noted by Schöck and Perrimon (47), this boundary is scalloped, pulling in toward the amnioserosa at AS-AS interfaces and thus implying forces at those locations that pull on the tissue boundary. Furthermore, throughout the course of germband retraction, the GB-AS boundary becomes more scalloped – i.e., the boundary cusps become sharper while the sections between become more highly curved. Qualitatively, the increased scalloping implies a weakening of tension in the GB-AS boundary relative to both AS-AS and GB-GB interfaces. To quantify these stage-dependent tension ratios, we measured triple-junction angles at the cusps and curvature of the boundary between cusps. Complete details and derivations of the geometry-tension relationships are provided in Supplemental Material. Results of the analysis are summarized in Fig. 1D, compiling data from nine embryos (with multiple time points for two embryos). This quantification confirms the qualitative assessment that the ratios γ_GB-AS_ / γ_AS-AS_ and γ_GB-AS_ / γ_GB-GB_ both decrease as retraction progresses.

To provide a smooth estimate for use in subsequent modeling, we fit each stage-dependent tension ratio to a sigmoidal function. These fits are shown in Fig. 1D: γ_GB-AS_ / γ_AS-AS_ is estimated as dropping from ∼2.4 to 0.6 with a sharp cross-over at 50% retraction (Hill coefficient ∼10); γ_GB-AS_ / γ_GB-GB_ similarly drops sharply from ∼17 to 5 at 42% retraction. Dividing these results, we estimate the ratio of amnioserosa to germband tension, γ_AS-AS_ / γ_GB-GB_ (dot-dash line in Fig. 1D), as increasing slightly at mid-retraction from ∼7 to 8. These time-dependent ratios place constraints on the tensions such that specifying any one of γ_AS-AS_, γ_GB-GB_ or γ_GB-AS_ implies specific values for the other two.

### Measuring retraction kinematics

Even with the above measurements, the model has some remaining free parameters that require target metrics for fitting the model to experiments. We thus quantify four metrics of retraction kinematics *in vivo.* To account for differences in developmental timing among embryos and experimental conditions, the raw time for our data and the data sets from Tomer *et al* (54) and Truong *et al* (55) are scaled by factors of 0.87, 0.75 and 1.5, respectively. This scaling aligns developmental progression temporally across all embryos analyzed.

The first two metrics are the positions of the **archicephalon** and **telson**, i.e., the dorsal-most points along the amnioserosa-head or amnioserosa-germband boundaries, respectively. In Fig. 1B, these are marked by magenta and green arrowheads. As retraction proceeds *in vivo,* the archicephalon remains nearly stationary while the telson moves away posteriorly. As shown in Fig. 1E, posterior movement of the telson accelerates through most of retraction and then slows as it approaches its final position at ∼85% of embryo length (*N* = 5 embryos). All measurements of archicephalon and telson positions are presented as relative to embryo length with 100% at the posterior end.

The third metric is amnioserosa cell shape as measured by the cellular aspect ratio distribution. To measure these shapes, we segmented time-lapse images of E-cadherin-GFP embryos *(N = 4* embryos, each with 49 to 104 measured cells). At the start of retraction, amnioserosa cells span lengths up to 20 times their width with a median aspect ratio, 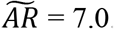. As highlighted in Fig. 1B, these cells bulge in-plane around their nucleus and extend long, protruding arms between neighboring cells. As retraction proceeds, amnioserosa cells round up to approximate regular polygons in plane. The resulting aspect ratio distributions are compiled in Fig. 1E (right). By the end of retraction, the distribution of aspect ratios has tightened considerably with the median aspect ratio just less than two.

The fourth and final metric is the stage-dependent shape of the GB-AS boundary. We will compare model results to previous experiments that quantified the changing shape of this contour as the germband uncurls (2).

### The model – constructing the initial configuration

The initial geometry of the model is matched to experimental images (Fig. 1B and 2A, t = 55 simulated minutes) and includes three cell types: isodiametric cells of both the head (Hd, magenta) and germband (GB, cyan); and highly elongated cells of the amnioserosa (AS, yellow, median aspect ratio of 7.7). Every cell in the amnioserosa is modeled explicitly, but the head and germband tissues are coarse-grained with each modeled “cell” representing ∼10 cells *in vivo.* To create the amnioserosa’s initial configuration, a 2D ellipse (4:3) was filled with 120 randomly distributed seeds that were spaced by Monte Carlo minimization of a 1/*r*^2^ repulsive interaction. These seeds were then used for a **Voronoi tessellation**. The tessellated ellipse was stretched to an aspect ratio of 12, giving the amnioserosa cells a median aspect ratio of 9.5, and then bent in the middle to form a U-shaped tissue (with an arc radius equal to 26% of embryo length). Each triple junction node of this U-shaped tissue (yellow in Fig. 2A) was then mapped onto the surface of a 3D prolate ellipsoid (1:1:2.5) that approximates the shape of a *Drosophila* embryo. This amnioserosa tissue shape corresponds to that observed *in vivo* nearly midway through retraction. Before that time, the amniosersoa is folded over itself on the dorsal surface and is not amenable to modeling as a single surface-wrapped monolayer. The first time point in model simulations is thus taken as 55 min after retraction commences.

Germband/head cells (cyan/magenta in Fig. 2A) were configured by seeding, spacing, and tessellating the ellipsoid’s surface using an ellipsoidal distance function,

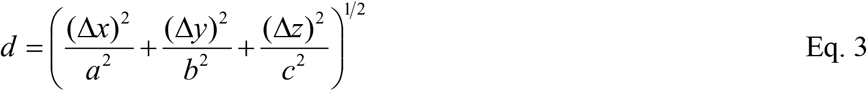

where *a* = *b* = 1 and *c* = 2.5 represent the semi-minor and semi-major radii corresponding to the *x, y,* and *z*-axes. Germband/head cells that lay completely within the region previously assigned to amnioserosa were discarded; those partially within that region were trimmed to form a continuous 2D mesh of connected polygonal cells with nodes constrained to lie on the surface of the 3D ellipsoid. The non-amnioserosa cells were assigned to head or germband depending on whether their centroids were anterior or posterior to the archicephalon (magenta arrowhead in Fig. 2A).

Experimental cell counts show that embryos typically contain about 120 amnioserosa cells and 3000 germband/head cells. The models presented here each use one of three initial configurations created as described above with 120 amnioserosa cells and approximately 300 germband/head cells. Models were also tested that eliminated the coarse graining and specified approximately 3000 germband/head cells. Simulations of germband retraction using the coarsegrained model took approximately 20 minutes (on a 3.40-GHz desktop PC); those using the denser mesh took approximately 4 hours. In representative simulations, results using the denser mesh matched those of the faster coarse-grained model.

### The model – matching in vivo kinematics using experimentally constrained parameters

Based on the experimental results above and some simplifications, the model’s nine potentially free parameters can be reduced to one free parameter ratio. First, the sigmoidal fits to the experimental tension ratios are used to lock the values of γ_GB-AS_ and γ_AS-AS_ to that of <γ_GB-GB_>. Since the model coarse-grains the germband with larger simulated cells, the tension ratios were scaled by the square root of 10. Second, the head region is covered by cells that closely resemble germband cells, but that move very little during retraction *in vivo* (Fig. 1A, B). The model limits movement of head cells without hindering movement of bordering germband cells by choosing larger values for head-region tensions: γ_Hd-Hd_ is taken as double <γ_GB-GB_>; γ_Hd-AS_ is taken as double γ_GB-AS_; and γ_GB-Hd_ is taken as equal to <γ_GB-GB_>. All six active edge tensions are thus defined in terms of the directionally-averaged and temporally varying GB-GB tension. The directional variation of γ_GB-GB_ in the model is then defined according to Eq. 2 with the experimentally determined polarization of germband cells implemented as *β*_0_ parallel to the closest section of GB-AS boundary and *α* set to 15%, which matches the largest observed value (and is slightly larger than the fitted values). With these relationships, nine potentially free parameters are reduced to two, <γ_GB-GB_> and the effective viscosity µ. In practice, only the ratio of these two parameters matters for the retraction kinematics, and that ratio is swept through a wide range of possible values using an optimizing **golden section search algorithm**. At each time step, this search determines the value of <γ_GB-GB_> / µ that yields a best match for the telson position. To yield smoother retraction profiles, the telson position target is taken from the quantic regression trendline of the average *in vivo* telson kymograph plots (black line, Fig. 1B). This procedure is replicated for three different initial cell meshes.

The results for one representative sample of the best fits are shown in Fig. 2 and Movie S2. In Fig. 2A, the final time point indicates a completely retracted germband. In Fig. 2B, the telson position completely overlaps the experimental target curve, just as the search was designed to accomplish. More importantly, the best fit based on telson positions also yields archicephalon positions and amnioserosa aspect ratios that are good matches to experimental observations. Furthermore, as shown in Fig. 2C, using the shape of the GB-AS boundary contour to estimate retraction progress following Lynch et al (2) yields results for the model that fall within experimental bounds. Interestingly, the best-fit tension set increases all tension values (relative to µ) in the latter half of retraction (Fig. 2D). This result reasonably explains the accelerating rate at which the telson moves posteriorly.

### Model comparisons to mutants and microsurgery experiments

Previous experiments have investigated the ability of germband retraction to proceed when the amnioserosa is compromised by mutation or microsurgery (1-3, 49, 52). We thus test whether the model reproduces these results using the same best fit parameters. To mimic cell death or ablation of amnioserosa cells *in silico,* the active edge tensions between dead/ablated cells are set to zero and those between living and dead/ablated cells are set to half their best fit value. All cells in the model, living or dead, retain their passive viscosity and incompressibility.

The first experimental perturbation reproduced with the model is ablating a small set of cells near the GB-AS boundary. As shown in Fig. 3A, ablating these cells *in silico* has little effect on retraction (see also Movie S3). This is exactly what was observed for *in vivo* laser microsurgery experiments (images from 2 reproduced in Fig. 5A).

**FIGURE 3.**
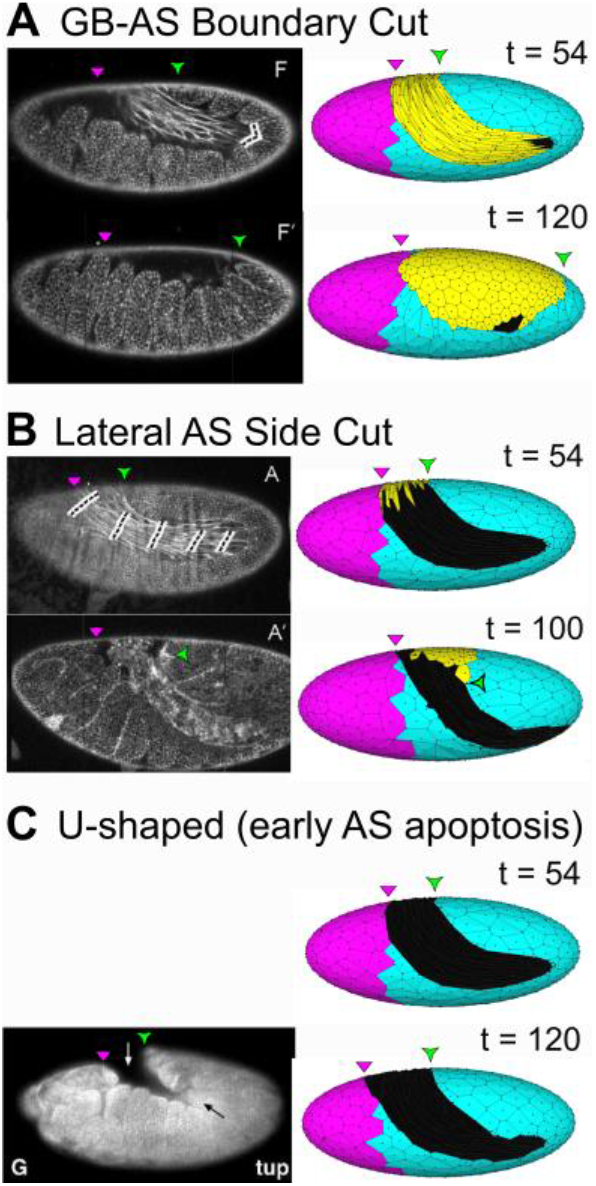
Replicating microsurgery results and mutant phenotypes. (A-B) Laser microsurgery experiments that ablated portions of the amnioserosa (images to left from 2; dashed lines mark laser targets) and their replication in silico using best fit model with γ_AS-AS_ = 0 for ablated cells (black). (C) U-shaped mutant tail-up after death of amnioserosa and failure of germband retraction (image to left from 3) with its replication in silico by assigning γ_AS-AS_ = 0 for all dead amnioserosa cells (black).

The second modeled perturbation is ablating half the amnioserosa, i.e., one of its lateral flanks. As shown in Fig. 3B, the remaining intact amnioserosa cells do round up (see also Movie S4). The unbalanced mechanical forces cause the posterior end of the germband to twist: the telson rotates toward the ‘wounded’ side and germband retraction fails. A similar result with a similarly twisted germband is observed when *in vivo* laser microsurgery is used to cut one lateral side of the amnioserosa (images from 2 reproduced in Fig. 5B).

The third modeled perturbation is complete removal of the amnioserosa. This happens in several U-shaped group mutants in which amnioserosa cells undergo premature apoptosis (3). As shown in Fig. 3C and Movie S5, *in silico* death of the entire amnioserosa yields only residual telson movement and a failure of retraction – resulting in a final morphology that resembles the eponymous morphology of the U-shaped mutant *tail up* (tup; image from 3 reproduced in Fig. 5C). All three of these *in silico* results match *in vivo* experiments without any modification of our best fit model save removing the edge tensions of dead/ablated amnioserosa cells.

As a final test, we assess the model’s ability to reproduce partial rescue of the U-shaped mutant *hindsight* (hnt) by overexpression of insulin receptor (inr, Fig. 4A reproduced from (1). We model *hnt* mutants by removing all amnioserosa cells, just as above, and then explore variations of germband tensions that could represent physical manifestation of the inr-rescue mechanism (any upstream signaling is not included in the model). Retraction defects were not rescued by increasing or decreasing germband tensions overall or by increasing the strength of their polarization (30% case shown in Fig. 4B). On the other hand, partial rescues were produced by simultaneously strengthening and orthogonally reorienting germband polarization. This rotated polarization yields its strongest germband tensions parallel to the GB-AS boundary, i.e., along the rostral-caudal axis. Minimal retraction was produced by doubling the reoriented polarization (Fig. 4C), but substantial rescues were produced for reoriented polarization with *α* = 80% or more (Fig. 4D and Movie S6). Clearly, the inr-rescue mechanism cannot simply involve strengthening or restoring a deficit of normal germband polarization, but seems to require its reorientation.

**FIGURE 4.**
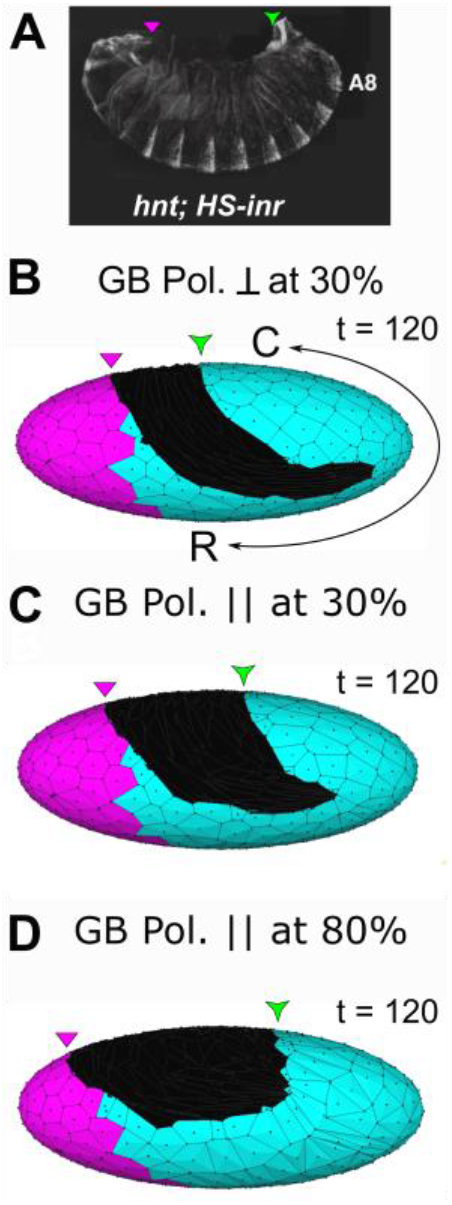
Rotated germband polarization yields partial rescue of retraction. (A) Partial rescue of U-shaped mutant hnt by ubiquitous overexpression of insulin receptor (image from 1). (B-D) In silico replication of rescue attempts showing U-shaped mutant simulations with modified germband polarization: (B) doubled (30%) with native orientation (⊥ to rostral-caudal axis); (C) doubled (30%) with rotated orientation (∥ to rostral-caudal axis); and (D) increased more than 5-fold (80%) with rotated orientation. Images show each model’s final morphology. R-C, rostral-caudal axis in (B) applies to all panels.

### Robustness with respect to the set of active tensions

Having replicated normal and aberrant germband retraction, we now explore its robustness and critical dependencies. We test the role of three key edge tensions: γ_AS-AS_, γ_GB-AS_, and γ_GB-GB_ – increasing each four-fold relative to the other two and labeling these tests “amnioserosa-dominant,” “boundary-dominant,” and “germband-dominant,” respectively (see Table S1 for a list of the specific sets of tensions). The amnioserosa-dominant test allows germband retraction to proceed to completion (Fig. 5A) and the boundary-dominant test yields nearly complete retraction, stopping at a telson position of 82% embryo length (Fig. 5B). On the other hand, the germband-dominant test fails to retract (Fig. 5C). The stiffer germband resists retraction until any asymmetry in the model’s Voronoi-tessellated cell configuration triggers an instability that causes the entire posterior three-quarters of the embryo to twist. This twisting is illustrated in the amnioserosa tissue sketch depicting how the tissue shifts from an initial U-shape to a curved pennant rather than a round ellipse (compare yellow sketches in Fig. 5 A-C).

Taking a cue from our simulations of genetic rescues above, we find that a germband-dominant case can successfully retract, but only with a greatly strengthened and rotated polarization (Fig. 5D). In the example shown for a rotated *α* = 80% case, the telson only retracts to 73% of embryo length, but the head is simultaneously pulled ventrally and the archicephalon moves anteriorly. The archicephalon-telson separation increases enough to say retraction succeeds, albeit in a manner that shifts the entire amnioserosa anteriorly.

**FIGURE 5.**
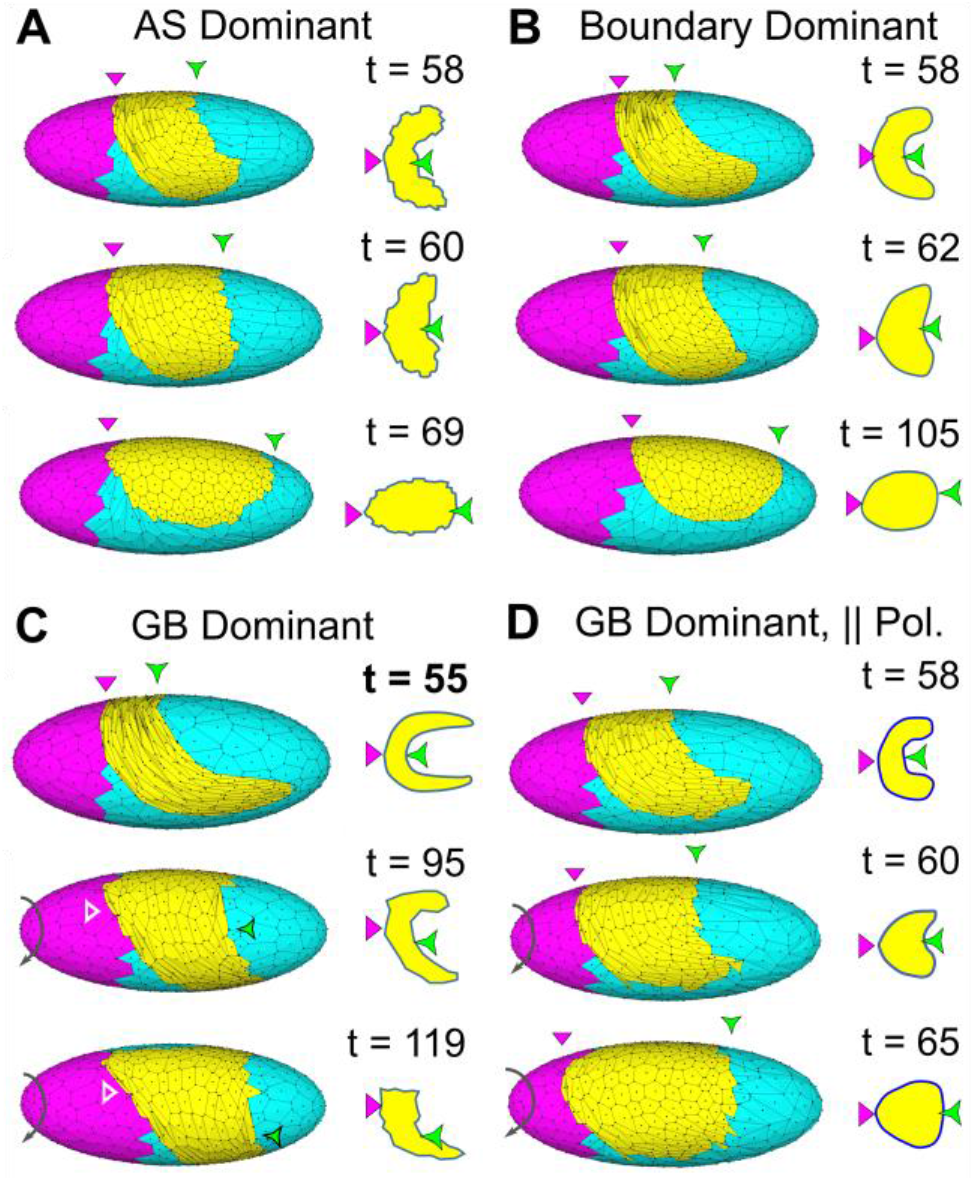
Retraction does not require fine-tuned tensions. (A-D) Models run with widely varying tension ratios (see Table S1 for details). Select time steps are shown with archicephalon/telson positions highlighted (magenta/green arrowheads). Corresponding amnioserosa tissue shapes are sketched at right. To better view amnioserosa movement and shape, model output images in (C-D) are rotated as indicated by gray arrows (less than ¼ turn). All models started from the same initial configuration at t = 55, but this configuration is only shown in (C).

These tests show that retraction requires either sufficient amnioserosa and boundary tensions to overcome resistance of the germband or a reoriented polarization in the germband that allows it to drive its own retraction. Otherwise, the process of germband retraction is quite robust to variations among the key active tensions.

### Critical role of elongated amnioserosa cells

Our best fit model and *in silico* mutants show that retraction is amnioserosa and boundary driven. On the other hand, retraction is quite robust to variations in γ_AS-AS_ and γ_GB-AS_, so what aspects of the amnioserosa are actually critical? To answer this question, we varied its initial cell shapes without changing its initial tissue shape. When individual amnioserosa cells started out almost round (median aspect ratio of 1.3), these models yielded essentially no retraction (Fig. 6A-B and Movie S7). Despite using the same best-fit tensions as above, the archicephalon and telson barely moved and the only discernable change with time was a roughening of the GB-AS boundary. Even when the tensions were modified to enhance the contribution of the boundary (γ_GB-AS_ increased 5-fold), simulations with isodiametric amnioserosa cells failed to retract (Fig. S2).

The critical role of initially elongated amnioserosa cells can be understood from a force perspective. The initiation of retraction requires a different balance of forces at different locations along the GB-AS boundary. Near the telson (Fig. 6C), the germband must overcome resistance from the amnioserosa to pull the telson posteriorly (to the right). Near the lateral tips of the amnioserosa (Fig. 6D), the amnioserosa must overcome resistance of the germband to move the boundary anteriorly (to the left). The key to satisfying both conditions is elongated amnioserosa cells. Their elongation near the telson is parallel to the tissue boundary, yielding a lower local density of boundary-intersecting AS-AS interfaces compared to GB-GB interfaces. In contrast, their elongation near the lateral tips is perpendicular to the tissue boundary, yielding a higher local density of boundary-intersecting AS-AS interfaces. With these configurations, both force balance conditions could be satisfied with a wide range of γ_GB-GB_ and γ_AS-AS_ values. If the amnioserosa cells began with nearly round shapes, then the relative density of AS-AS and GB-GB interfaces would be similar at both locations along the boundary and no choice of uniform γ_GB-GB_ and γ_AS-AS_ values could satisfy both force balance conditions. If amnioserosa cells began with slightly elongated shapes, then retraction could initiate and proceed until the amnioserosa cells became sufficiently round and the different force balance conditions could no longer be satisfied. In tests with intermediate initial cell elongations, the model yields exactly this sort of partial retraction.

**FIGURE 6.**
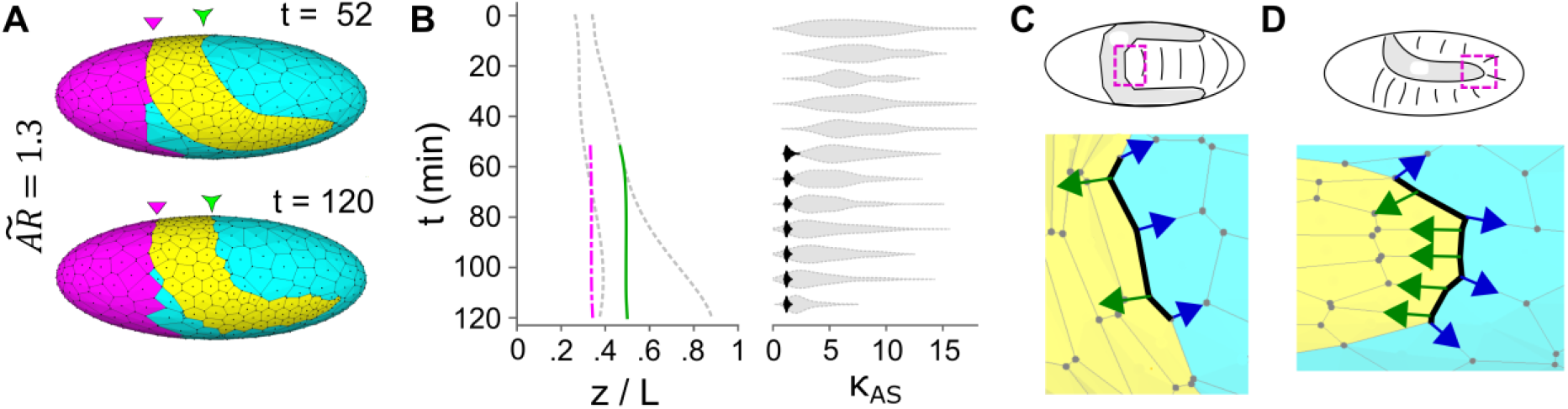
Initial amnioserosa cell shape determines the extent of germband retraction. (A) Model morphology and (B) retraction kinematics when amnioserosa cells begin as nearly isodiametric. 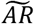 is initial median cellular aspect ratio. (C-D) Close-up views of a mesh as indicated by magenta dashed outlines on embryo sketches. Views taken early in retraction (t = 63) when amnioserosa cells begin with their normally elongated shapes: amnioserosa cells (yellow) and germband cells (cyan). Arrows indicate where tensions along AS-AS (green) and GB-GB (blue) interfaces pull on a segment of the GB-AS boundary (black line).

## DISCUSSION

The finite element model presented here uses 2D cells confined to the surface of a 3D last. This configuration models epithelial mechanics in *Drosophila* embryos under appropriate ellipsoidal boundary conditions – translating cell-level forces into embryo-spanning tissue interactions. The model successfully reproduces the process of germband retraction, including wild-type retraction kinematics, select mutant phenotypes, and the results of microsurgical interventions.

The cell-level forces used in the model are active tensions along cell-cell interfaces. Time-dependent ratios of these tensions were inferred from experimental observations of cell morphology. Most notably, increased scalloping of the lateral GB-AS boundary halfway through retraction implies a relative weakening of tension along the tissue boundary. Once the complete set of tensions is estimated by matching modeled and observed rates of retraction – varying just one remaining free parameter – it becomes evident that all of the active tensions increase throughout retraction. These increases are needed to match the observed accelerating rate of telson movement. The relative weakening of the boundary results because the mid-retraction transition to higher tensions is stronger within each tissue (γ_GB-GB_ and γ_AS-AS_) than along the boundary (γ_GB-AS_). The signal that triggers these changes in active tensions is yet unknown, but one interesting possibility is an observed mid-retraction pulse of the insect hormone ecdysone (67).

Several previous studies have shown that the amnioserosa plays a key role in germband retraction (1-3, 47-52). Our best fit model similarly suggests that retraction is an amnioserosa-driven process. Nonetheless, others present evidence that the germband actively contributes and may in certain circumstances drive its own retraction (1, 48, 52). The only mechanism by which this could occur in our model is via anisotropy among germband cell-edge tensions. Previous modeling and experiments have suggested that such anisotropy is present (15, 48, 63), but our analysis of germband cell morphology shows that this anisotropy is actually oriented to resist retraction. In attempting to simulate the mutant rescue, *hnt; HS-inr* (1), in which amnioserosa cells die prematurely, we generate a partial rescue by rotating a strengthened germband anisotropy by 90°. This is an interesting hypothesis, but alternative mechanisms remain for germband contributions that are beyond the scope of the current model, for example, the deepening of germband segmental grooves or the orientation of microtubules to provide anisotropic cell interiors, as is known to occur later in dorsal closure (68).

The results presented here also inform the explanation of other mutant phenotypes and perturbations in which the amnioserosa’s structural integrity is compromised. For example, germband retraction fails after gastrulation-stage heat shocks that lead to gaping holes in the amnioserosa (69). It also fails in the mutants *scarface* (70) and *myospheroid* (71), which have compromised adhesion between amnioserosa cells and their basement membrane. Interestingly, *myospheroid* mutants often twist (71). The model reproduces such twisting when amnioserosa tension is asymmetrically compromised (Fig. 3B) or just too weak (Fig. 5C). For the models used here, the absence of living amnioserosa cells is the same as the absence of amnioserosa cell-edge tensions. The same models used to recapitulate U-shaped mutants could thus also describe retraction failures in amnioserosa-specific knockdown of RhoA (dominant negative UAS-rho1^N17^:c381-Gal4; (47)). RhoA is an upstream regulator of myosin II, so expression of its dominant negative form inhibits tension production via non-muscle myosin II. Our model predicts a failure of germband retraction in any genetic or environmental background that disrupts the structural integrity and/or force production within the amnioserosa.

Although we have found specific and interesting relationships among the active tensions driving germband retraction, these tensions can vary widely while still yielding retraction, albeit with different kinematics. Retraction in the model is thus robust with respect to fine-tuning of the cellular tensions. Instead, the primary factor controlling the extent of retraction *in silico* is the initial shape of amnioserosa cells. If these cells are sufficiently elongated at the start, then retraction will eventually succeed for a wide range of active tensions – failing only if the germband resists deformation too strongly. On the other hand, if amnioserosa cells are close to isodiametric at the start, then retraction fails. The embryo-spanning process of uncurling of the germband requires an interesting configuration of tissue-level forces: the amnioserosa must pull harder than the germband on their lateral interfaces, but the germband must pull harder at the tissues’ dorsal interface. Isodiametric amnioserosa cells cannot simultaneously satisfy both requirements.

The elongated shapes of amnioserosa cells are specified as initial conditions in the model, but these shapes are established in an embryo during the previous process of germband extension. The model thus reveals a key link by which one morphogenetic event sets the stage for the next. During germband extension, the germband extends via convergent cell neighbor exchanges (14, 63). These tissue movements guide amnioserosa cell elongation, which is actively driven by anisotropic trafficking of cell adhesion molecules and microtubule rotation/extension (16, 24). At a cellular level, this elongation is dissipative: amnioserosa cells are not stretched elastically and do not store potential energy (72). Nonetheless, because these cells elongate without exchanging neighbors (16, 24), the amnioserosa stores information. When active tensions increase during germband retraction, those in the amnioserosa drive its cells back to the nearly isodiametric shapes they held before the paired processes began. Since amnioserosa cells are still connected to the same neighbors, this rounding up drives the tissue back to its initial dorsal location and shape, allowing it to assist in uncurling the germband.

Morphogenesis involves collective cell migration, which emerges from cell coordination at the tissue scale. Previous modeling efforts have sought to identify the mechanism(s) that drive such migration, i.e., which cells and even which subcellular components create the forces necessary to effect morphological change (e.g., 4, 5, 8, 37, 48). For the models of germband retraction investigated here, morphogenesis is not strongly dependent on the specific allocation of cellular tensions. This observation begs a question: although force generation is certainly necessary, might the sources and magnitudes of the forces be less important, or at least no more important, than their coordination by sources of morphological information? In the case of germband retraction, we have found that the key sources of morphological information are amnioserosa cell geometry and maintenance of cell-cell connections. The cell geometry allows forces generated in individual amnioserosa cells to yield mesoscopic tensions that pull in appropriate and yet different directions at different locations. The conserved connection topology allows amnioserosa cell shapes established during germband extension to later robustly guide cells through germband retraction as they are reversed. It will be interesting to see if the storage of morphological information in previously deformed tissues proves to be a general principle of morphogenetic mechanics.

## AUTHOR CONTRIBUTIONS

G.W.B. and M.S.H. designed research; W.T.M., M.E.B. and M.S.H. performed and analyzed experiments; W.T.M., M.S.H, J.V. and G.W.B. did mathematical modeling; and W.T.M., M.S.H, J.V. and G.W.B. wrote the article.

## ACKNOWLEDGEMENTS

This work supported by the National Institutes of Health through Grant 1R01GM099107 and by the National Science Foundation through Graduate Research Fellowships to W.T.M. and M.E.B. The simulation viewer ChiChi3D was developed by Gregory Bootsma under Natural Sciences and Engineering Research Council (NSERC) funding to G.W.B.

