## Supplemental Material for "Elongated cells drive morphogenesis in a surface-wrapped finite element model of germband retraction"

### TABLES

| Description | $\gamma_{GB\text{-}GB}$ | $\gamma_{GB\text{-}AS}$ | $\gamma_{AS\text{-}AS}$ | $\gamma_{Hd\text{-}GB}$ | $\gamma_{Hd\text{-}AS}$ | $\gamma_{Hd\text{-}Hd}$ | Figure |
|---|---|---|---|---|---|---|---|
| Best Fit to GB Retraction[b] | 48(⊥), 36(‖)[a] | 71 | 118 | 42 | 143 | 84 | 2 |
| Microsurgery[c] | 48(⊥), 36(‖)[a] | 71 | 118 | 42 | 143 | 84 | 3A-B |
| (ablated AS) | | **(36)** | **(0 or 59)[d]** | | **(72)** | | |
| U-shaped mutants [c] | 48(⊥), 36(‖)[a] | 71 | **0** | 42 | 143 | 84 | 3C |
| Polarized rescues [c] | | | | | | | |
| ‖ 30% Polarized $\gamma_{GB\text{-}GB}$ | **55(⊥), 29(‖)[a]** | 71 | **0** | 42 | 143 | 84 | 4A |
| ⊥ 30% Polarized $\gamma_{GB\text{-}GB}$ | **29(⊥), 55(‖)[a]** | 71 | **0** | 42 | 143 | 84 | 4B |
| ⊥ 80% Polarized $\gamma_{GB\text{-}GB}$ | **8(⊥), 75(‖)[a]** | 71 | **0** | 42 | 143 | 84 | 4C |
| ↑ $\gamma_{GB\text{-}GB}$ by 4x | 40 | 20[e] | 10 | 40 | 40 | 80 | 5A |
| ↑ $\gamma_{AS\text{-}AS}$ by 4x | 10 | 20[e] | 40 | 10 | 40 | 20 | 5B |
| ↑ $\gamma_{GB\text{-}AS}$ by 4x | 10 | 40 | 10 | 10 | 40 | 20 | 5C |
| 80% Polarized $\gamma_{GB\text{-}GB}$ | 8(⊥), 72(‖)[a] | 20 | 10 | 40 | 40 | 80 | 5D |
| Less AS cell elongation[f] | 61(⊥), 45(‖)[a] | 90 | 150 | 53 | 180 | 106 | 6A-B |
| Attempted rescue[c,f] | 61(⊥), 45(‖)[a] | **450** | 150 | 53 | 180 | 106 | S2 |

[a]Polarized GB cells have maximum and minimum tensions perpendicular (⊥) and parallel (‖) to the nearest GB-AS boundary segment.

[b]Best fit parameters were found as described in text. Maximum values are listed for one mesh. See Fig. 2D for tensions through time.

[c]Best fit parameters were used with adjustments highlighted in **bold**. See text for details.

[d]Tension is zero between two ablated cells and halved along the interface between an ablated and normal cell.

[e] The GB-AS boundary tension, $\gamma_{GB\text{-}AS}$, is doubled to prevent topological complications due to the surface constraint of the last.

[f]Maximum best fit parameters used from different original mesh to better align with initial conditions of special AS meshes.

**TABLE S1. Tension Parameters for Model Variants**. Each cell-cell interface is assigned a line tension determined by the two adjacent cell types: GB = germband; AS = amniosersosa; Hd = head.

### MOVIES

**MOVIE S1. *In vivo* timelapse video of germband retraction.** Corresponds to Fig. 1B. Viewed dorsolaterally. Time is scaled to match plots in Fig. 1E.

**MOVIE S2. Best-fit simulation**. Corresponding to Fig. 2A.

**MOVIE S3. *In silico* completion of germband retraction following minor ablation.** Replicates two-cell ablation due to microsurgery. Corresponds to Fig. 3A.

**MOVIE S4. *In silico* twisting of germband retraction after lateral amnioserosa ablation.** In tact far lateral side induces germband twist. Corresponds to Fig. 3B.

**MOVIE S5. *In silico* failure of germband retraction in U-shaped mutant replicate.** All amnioserosa cell tensions are removed as per the text. Corresponds to Fig. 3C.

**MOVIE S6. *In silico* reoriented polarization partially rescues U-shaped mutant.** Replicates inr-rescue using a rotated 80% germband cell polarization. Corresponds to Fig. 4D.

**MOVIE S7. Germband retraction dependends on amnioserosa cell shape.** Reduced cell aspect ratio results in failure of retraction (see text for details). Corresponds to Fig. 6A.

### SUPPORTING RESULTS AND DISCUSSION

#### *Estimating Relative Tensions from Images of the GB-AS Boundary*

The process of germband retraction in *Drosophila* embryos involves two connected epithelia, amnioserosa (AS) and germband (GB). Fig. S1 below shows the cellular and tissue geometry of the GB-AS boundary on the lateral flank of an embryo during late GB retraction. At this stage, the boundary typically has a scalloped appearance. As noted by Schöck and Perrimon (1), this scalloped appearance indicates that AS-AS interfaces are pulling on the attached GB (yellow arrowheads in Fig. S1). By careful measurements of the GB-AS boundary, one can estimate relative tensions along three types of cell-cell interfaces: AS-AS, GB-GB and GB-AS.

The first key measurements are the angles at triple junctions where AS-AS interfaces intersect the GB-AS boundary. An example is shown as the inset expanded in Fig. S1A. The relationships among the interfacial tensions ($\gamma$'s) can be found by considering force balance under conditions very close to static equilibrium. If the *x*-axis is taken as parallel to tension in the AS-AS interface ($\gamma_2$), then force balance yields:

$$F_{Net,x} = 0 = \gamma_2 - \gamma_1 \cos(180° - \theta_3) - \gamma_3 \cos(180° - \theta_1) \quad \text{Eq. S1}$$

$$F_{Net,y} = 0 = \gamma_1 \sin(180° - \theta_3) - \gamma_3 \sin(180° - \theta_1) \quad \text{Eq. S2}$$

which can be solved for the tension ratios

$$\gamma_1/\gamma_2 = \frac{-\sin\theta_1}{\sin(\theta_1+\theta_3)} \quad \text{and} \quad \gamma_3/\gamma_2 = \frac{-\sin\theta_3}{\sin(\theta_1+\theta_3)} \quad \text{Eq. S3-4}$$

Both are ratios of tension in the GB-AS boundary to tension along AS-AS interfaces. If the boundary is pulled tauter, $\theta_1$ and $\theta_3$ decrease (approaching 90°), and the ratios above increase. If the amnioserosa pulls more strongly, the ratios decrease because $\theta_2$ becomes sharper while $\theta_1$ and $\theta_3$ become larger. The ratios become less than one when $\theta_1$ and $\theta_3$ are greater than 120°. For a given embryo, we calculate the mean $\gamma_{GB\text{-}AS}/\gamma_{AS\text{-}AS}$ by averaging both ratios over multiple triple

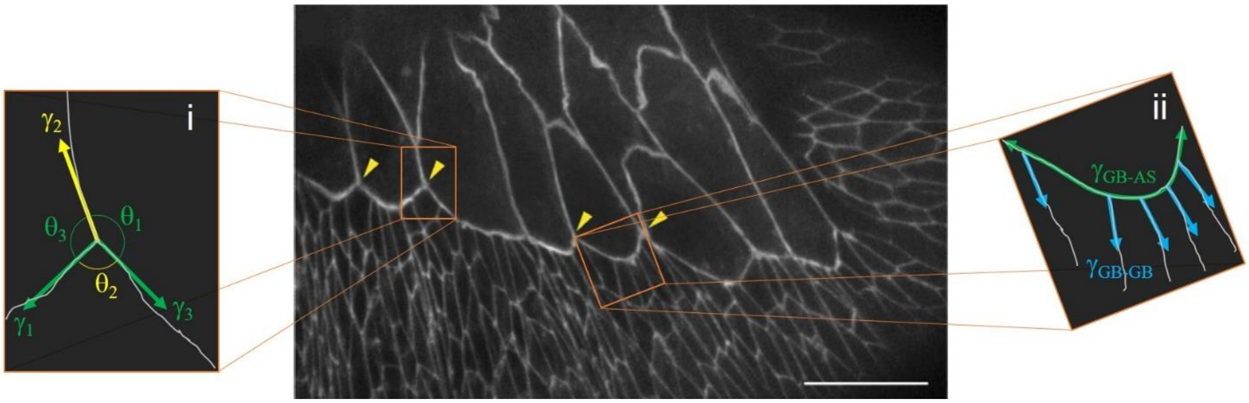


**FIGURE S1.** Geometry of the GB-AS boundary during late germband retraction. Confocal image of an E-cadherin-GFP labeled embryo is reproduced from Schöck and Perrimon (1). Large cells at top left are amnioserosa; smaller cells at bottom and right are germband. Scale bar is 20 µm. (i) Geometry of a triple-junction at which an AS-AS cell interface, carrying tension $\gamma_2$, intersects the GB-AS boundary, carrying tensions $\gamma_1$ and $\gamma_3$ to left and right respectively. (ii) Curved section of the GB-AS boundary between two AS-AS cell interfaces. This section of interface carries average tension $\gamma_{GB\text{-}AS}$ and is intersected by five GB-GB cell interfaces, carrying average tension $\gamma_{GB\text{-}GB}$.

junctions. These means are plotted in Fig. 1D of the main text. For the embryo shown in Fig. S1, these calculations yield a mean $\gamma_{\text{GB-AS}}/\gamma_{\text{AS-AS}}$ ratio of ~ 0.9.

The second key measurement is the curvature of GB-AS boundary sections between AS-AS interfaces. An example is shown as the inset expanded in Fig. S1C. These sections can be analyzed with a 2D analogy to a Laplace pressure that relates the radius of curvature ($R_c$) and tension of the boundary ($\gamma_{\text{GB-AS}}$) to the difference in force per unit length acting normal to the boundary ($\Delta F/L$):

$$\frac{\Delta F}{L} = \frac{\gamma_{\text{GB-AS}}}{R_c} \qquad \text{Eq. S5}$$

We approximate the forces as being carried only by cortical tensions along cell-cell interfaces. The force difference on a boundary section between AS-AS interfaces is thus the tension applied by all the intersecting GB-GB interfaces. Given that these interfaces are roughly normal to the boundary, the force difference is approximated by counting intersections, $\Delta F = N \langle\gamma_{\text{GB-GB}}\rangle$. Substituting the force difference into Eq. S5 and rearranging yields the tension ratio

$$\frac{\gamma_{\text{GB-AS}}}{\langle\gamma_{\text{GB-GB}}\rangle} = \frac{NR_c}{L} \qquad \text{Eq. S6}$$

If the boundary tension $\gamma_{\text{GB-AS}}$ increases, the increase in tension ratio is evidenced by a flattening of the boundary (larger $R_c$). If the germband pulls more strongly or the boundary weakens, the tension ratio decreases because the boundary becomes more highly curved (lower $R_c$). As above, we calculate a mean $\gamma_{\text{GB-AS}}/\gamma_{\text{GB-GB}}$ for each embryo by averaging over multiple arc-like boundary sections. These means are plotted in Fig. 1D of the main text. For the embryo shown in Fig. S1, these calculations yield a mean $\gamma_{\text{GB-AS}}/\gamma_{\text{GB-GB}}$ ratio of ~ 5.

Note that a close inspection of the GB-AS boundary shows that it is bent slightly at each intersection with a GB-GB interface. Using a triple-junction analysis similar to that above, one can estimate that a tension ratio of ~ 3 would bend the boundary at each GB-GB interface by less than 10°. Given the short lengths of GB-AS boundary between each GB-GB interface, it is difficult to estimate angles. Estimating curvature is thus a means to average over the cumulative bending by a succession of GB-GB interfaces. The radius of curvature for each GB-AS boundary section was estimated visually in ImageJ by overlaying an adjustable arc on the image.

*Rescuing retraction when amnioserosa cells are initially isodiametric*

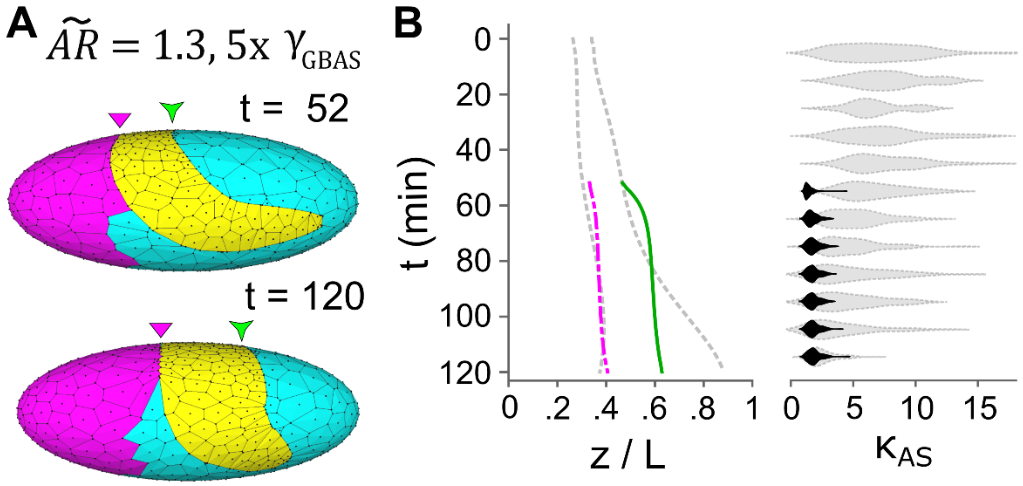


**FIGURE S2**: The failure of germband retraction with initially rounded amnioserosa cells (Fig. 6A-B of main text) can be partially rescued by dramatically increasing tension along the GB-AS border (5x $\gamma_{GB\text{-}AS}$). (A) Initial and final time points show extent of retraction. (B) Retraction kinematics as in previous figures: curves track archicephalon and telson movements as a fraction of embryo length (z / L) for the model (dot-dashed magenta and solid green respectively) versus *in vivo* data (dashed gray; smooth quantic regressions shown); distributions are for amnioserosa cell aspect ratios ($\kappa_{AS}$): model (black) versus *in vivo* data (light gray).

### GLOSSARY OF TERMS

The following glossary describes how terms are used specifically within the context of mechanical models for *Drosophila* development.

**Amnioserosa (AS)**: an extraembryonic epithelioid tissue that covers parts of the dorsal surface of *Drosophila* embryos during mid embryogenesis (gastrulation to dorsal closure)

**Archicephalon**: location within a *Drosophila* embryo corresponding to the anterior-most point along the boundary between the head and dorsal amnioserosa

**Germband (GB)**: embryonic tissue that covers the ventral and lateral surface of insect embryos and develops into the segmented larval ectoderm; in *Drosophila*, the germband has 12 segments: T1-T3 and A1-A9.

**Golden section search algorithm**: method for searching a parameter range for an extremum of a function (here used to find model parameters that minimize the difference between modeled and measured telson positions); as the search algorithm proceeds, narrower ranges of parameters are tested using golden ratios to make successive parameter guesses

**Isodiametric**: describing a shape having similar spans when measured in any direction; for 2D epithelial cells, this implies shapes that are roughly regular convex polygons

**Kinematics**: description of cell and tissue movements and deformations, *e.g.*, telson velocity or rate of change in amnioserosa cell aspect ratio

**Last**: a rigid mechanical form over which a surface is wrapped; term commonly used by cobblers in describing the foot-shaped form over which materials are wrapped to make shoes

**Reynolds Number**: *dimensionless* ratio of inertial to viscous forces in fluid flow; low Reynolds numbers are prevalent in morphogenesis where inertia has negligible contribution (2)

**Rostral-caudal axis**: axis lying along the length of the germband; at the beginning of germband retraction, the axis wraps around the posterior end of the embryo with its rostral or nose end located anterioventrally and its caudal or tail end located anteriodorsally

**Telson**: location within a *Drosophila* embryo corresponding to the anterior-most point along the boundary between the dorsal amnioserosa and the caudal tip of the germband

**Voronoi tessellation**: method of dividing a 2D space completely into polygonal regions (cells) based on a finite number of seed points; all points within a polygon share the same closest seed
